# *Single-cell transcriptome dataset of human and mouse* in vitro *adipogenesis models*

**DOI:** 10.1101/2023.03.27.534456

**Authors:** Jiehan Li, Christopher Jin, Stefan Gustafsson, Abhiram Rao, Martin Wabitsch, Chong Y. Park, Thomas Quertermous, Ewa Bielczyk-Maczynska, Joshua W. Knowles

**Affiliations:** Division of Cardiovascular Medicine, Department of Medicine, Stanford University School of Medicine, Stanford, CA, 94305, USA; Stanford Diabetes Research Center, Stanford University School of Medicine, Stanford, CA, 94305, USA; Stanford Cardiovascular Institute, Stanford University School of Medicine, CA, 94305, USA; Clinical Epidemiology Unit, Department of Medical Sciences, Uppsala University, Uppsala, Sweden; Department of Bioengineering, Stanford University, Stanford, CA 94305, USA; Department of Pediatrics and Adolescent Medicine, Center for Rare Endocrine Diseases, Division of Pediatric Endocrinology and Diabetes, Ulm University Medical Centre, Ulm, 89075, Germany; Stanford Prevention Research Center, Stanford University School of Medicine, Stanford, CA, 94305, USA

## Abstract

Adipogenesis is a process in which fat-specific progenitor cells (preadipocytes) differentiate into adipocytes that carry out the key metabolic functions of the adipose tissue, including glucose uptake, energy storage, and adipokine secretion. Several cell lines are routinely used to study the molecular regulation of adipogenesis, in particular the immortalized mouse 3T3-L1 line and the primary human Simpson-Golabi-Behmel syndrome (SGBS) line. However, the cell-to-cell variability of transcriptional changes prior to and during adipogenesis in these models is not well understood. Here, we present a single-cell RNA-Sequencing (scRNA-Seq) dataset collected before and during adipogenic differentiation of 3T3-L1 and SGBS cells. To minimize the effects of experimental variation, we mixed 3T3-L1 and SGBS cells and used computational analysis to demultiplex transcriptomes of mouse and human cells. In both models, adipogenesis results in the appearance of three cell clusters, corresponding to preadipocytes, early and mature adipocytes. These data provide a groundwork for comparative studies on human and mouse adipogenesis, as well as on cell-to-cell variability in gene expression during this process.

## Background & Summary

Adipose tissue carries out multiple roles that affect whole-body metabolism. In addition to storing energy in the form of lipids, it contributes to the homeostatic maintenance of blood glucose levels by taking up glucose in response to insulin and regulates the function of other metabolic organs by secreting hormones such as leptin and adiponectin^1,2^.

Adipogenesis is a differentiation process in which fat-specific progenitor cells (preadipocytes) convert into adipocytes, which carry out key metabolic functions of the adipose tissue. *In vivo*, preadipocytes are located in proximity of blood vessels within adipose tissue and contribute to adipose tissue maintenance and expansion in obesity^3^. Dysregulation of adipogenesis can result in metabolic disease, including insulin resistance and type 2 diabetes.^4^

Several preadipocyte *in vitro* models are routinely used to study the molecular regulation of adipogenesis. The most commonly used *in vitro* models include the immortalized mouse 3T3-L1 cell line^5^ and the primary, non-immortalized, non-transformed human Simpson-Golabi Behmel syndrome (SGBS) cell line^6^. These cellular models brought on major breakthroughs in our understanding of molecular mechanisms of adipogenic differentiation, both in development and in obesity^7,8^. However, adipogenic models show high levels of cell-to-cell heterogeneity in their differentiation responses to stimuli^9^. This heterogeneity can be due to multiple factors, including variations in preadipocyte commitment and stochasticity of responses to differentiation stimuli. Despite that, adipogenesis is often studied using bulk approaches, such as bulk RNA-Sequencing, which ignore the variability between individual cells, likely masking the presence of distinct cell subpopulations during adipogenesis.

Here, we present a single-cell RNA-Sequencing (scRNA-Seq) dataset collected before and during adipogenic differentiation of 3T3-L1 and SGBS cells to allow for analyses of heterogeneity of transcriptional states before and during adipogenesis, as well as comparisons between mouse and human models of adipogenesis. To minimize technical variation, at two time points (before and during adipogenic differentiation) mouse and human cells were mixed in equal ratios and subjected to scRNA-Seq, followed by computational demultiplexing and separation of data from mouse and human cells (**Fig. 1**). Through technical validation, we demonstrate quality of this dataset. By unsupervised clustering we identify cell populations that correspond to preadipocytes, differentiating and mature adipocytes in both models.

**Fig. 1.**
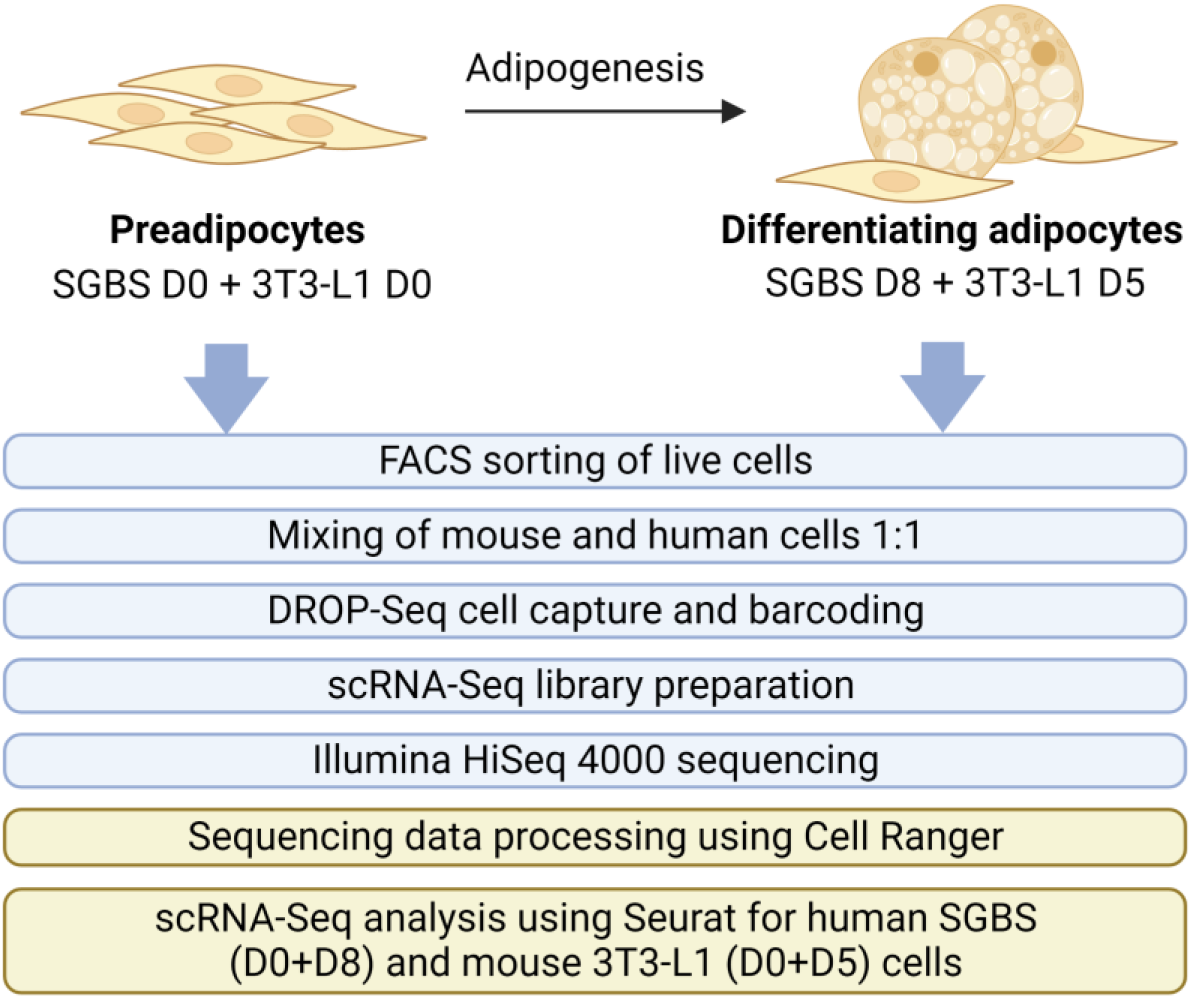
Workflow of scRNA-Seq of mouse and human adipogenesis. Human SGBS and mouse 3T3-L1 cells were analyzed at two time points, corresponding to before (D0) and during (D5 for 3T3-L1, D8 for SGBS) adipogenesis. At each time point, live cells were purified using exclusion of propidium iodide-stained cells by FACS. Equal numbers of SGBS and 3T3-L1 cells were then mixed, and subjected to microfluidic single-cell capture with GelBeads-in-emulsion (GEMs) using 10X Chromium Controller. Single-cell cDNA libraries were prepared using the Chromium Single Cell 3’ Library & Gel Bead Kit (10X Genomics), followed by sequencing on Illumina HiSeq4000. Computational analysis involved barcode processing, UMI counting, demultiplexing, gene and cell filtering, normalization, and clustering.

This dataset complements recent advances in characterizing the transcriptome of adipose tissue in human and mice at a single-cell^10–13^ and single-nucleus level^14^.

## Methods

### Cell culture

The 3T3-L1 preadipocyte cell line was maintained in Dulbecco’s Modified Eagle’s Medium (DMEM, Thermo Fisher) with 10% Fetal Bovine Serum (GeminiBio), 100 units/ml penicillin and 100 μg/ml streptomycin, in a humidified 5% CO2 incubator. For adipogenic differentiation cells were grown to confluency. 48 h past confluency, at day 0 of differentiation, cells were stimulated with 1 μM dexamethasone, 0.5 mM IBMX, 10 μg/ml insulin in growth medium. After 48 h the medium was changed to growth medium with 10 μg/ml insulin in growth medium until day 5.

The SGBS cell line was cultured and differentiated as previously described^6^. Cells were maintained in a humidified chamber at 37°C with 5% CO_2_, and the media was replaced every 2-3 days. The standard culture media used was composed of DMEM/Nutrient Mix F-12 (Invitrogen), supplemented with 33 uM biotin, 17 uM pantothenic acid, 10 % FBS and antibiotics (100 IU/ml penicillin and 100 ug/ml streptomycin). Differentiation was induced on D0, three days post-confluence, by the change of culture media to DMEM/F-12, 33 uM biotin, 17 uM pantothenic acid, 0.01 mg/ml human transferrin, 100 nM cortisol, 200 pM triiodothyronine, 20 nM human insulin (Sigma-Aldrich), 25 nM dexamethasone, 250 uM IBMX, 2 uM rosiglitazone, and antibiotics. After four days of differentiation, the medium was replaced with DMEM/F-12, 33 uM biotin, 17 uM pantothenic acid, 0.01 mg/ml human transferrin, 100 nM cortisol, 200 pM triiodothyronine, 20 nM human insulin and antibiotics. SGBS cells were cultured for eight days after the induction of differentiation.

### Single-cell sorting and cDNA library preparation

On the day of collection, cells were detached from culture plates using TrypLE Select Enzyme (Gibco), centrifuged at 300 x g for 5 min and resuspended in PBS with 0.04% Bovine Serum Albumin. Lack of staining with propidium iodine (PI) was used to sort live cells using Influx sorter (Beckman Dickinson). Equal numbers of SGBS and 3T3-L1 cells were mixed and subjected to single-cell capture on the 10X Chromium Controller device at Stanford Genomics Service Center during which single cells were encapsulated with individual Gel Beads-in-emulsion (GEMs) using the Chromium Single Cell 3’ Library & Gel Bead Kit (10X Genomics). In-drop reverse transcription and cDNA amplification was conducted according to the manufacturer’s protocol to construct expression libraries. Library size was checked using Agilent Bioanalyzer 2100 at the Stanford Genomics facility. The libraries were sequenced using Illumina HiSeq 4000.

### Raw data processing

Cell Ranger v2.10 was used for processing and analysing the raw single cell FASTQ files. The following genome builds were used: mm10 for the mouse genome, hg19 for the human genome. Quality control (QC) steps taken to assess the quality of the sequencing data and identify potential included: sample demultiplexing, read alignment and filtering, gene expression quantification, cell filtering and QC metrics, and data normalization and batch correction. Only reads mapping to mm10 or hg19 were used for downstream processing.

### Bioinformatic analysis of scRNA-Seq data

Seurat v4.3^15^ was used to merge processed data for two single cell sequencing runs, combining sequencing data from different stages of adipocyte differentiation. The data was first split between human and mouse data, pre-processed using Seurat, then log normalized. The major variable features within the processed data were identified using Variance Stabilizing Transformation. The gene matrix was then visualized and analysed using principal component analysis (PCA), with gene associations to each principal component displayed. Seurat’s *FindNeighbors* and *FindClusters* functions (resolution = 0.09) were used to identify groups within the samples. The data was further visualized via the PCA, Uniform Manifold Approximation and Projection (UMAP), and t-distributed Stochastic Neighbor Embedding (t-SNE) dimensional reduction techniques. Seurat’s *FindAllMarkers* function identified specific genes specific to each cluster, with previous annotations indicating that genes were clustered by stages in cell differentiation. Feature plots for specific differentiation features were visualized in a t-SNE plot and through heatmaps for each cluster using Seurat’s *DoHeatMap* and *FeaturePlot* functions.

## Data Records

Sequencing data have been submitted to the NCBI Gene Expression Omnibus (GSE226365). The dataset consists of raw sequencing data in FASTQ format, separated by the time point: D0 3T3-L1 and D0 SGBS (GSM7073976) and D5 3T3-L1 and D8 SGBS (GSM7073977). In addition, we provide processed data, separated by time point and cell line, including *barcodes.tsv, genes.tsv* and *matrix.mtx* files, listing raw UMI counts for each gene (feature) in each cell (barcode) in a sparse matrix format.

## Technical Validation

To validate the quality of our data, we investigated the technical quality control and the unsupervised clustering and its reproducibility between the two datasets.

### Quality control of the scRNA-Seq dataset

Interpretation of single-cell transcriptomics data is highly sensitive to technical artifacts. Sequencing data alignment using Cell Ranger led to the identification of comparable numbers of human and mouse cells within each of the analysed time points, as expected **(Table 1)**. We used further steps to filter cells, removing any multiplets and cells with fewer than 200 genes detected **(Fig. 2, Table 2)**.

**Table 1.** Detailed QC report of 10X Genomics sequencing files (Cell Ranger).

| Raw sequencing sample | SGBS D0, 3T3-L1 D0 |  | SGBS D8, 3T3-L1 D5 |  |
| --- | --- | --- | --- | --- |
| Number of reads | 320,829,287 |  | 334,091,518 |  |
| Q30 bases in barcodes | 96.9% |  | 97.5% |  |
| Q30 bases in RNA reads | 76.7% |  | 77.4% |  |
| Q30 bases in UMI reads | 96.8% |  | 97.6% |  |
| Mean reads per cell | 31,460 |  | 49,239 |  |
| Processed sample | SGBS D0 | 3T3-L1 D0 | SGBS D8 | 3T3-L1 D5 |
| Reads mapped to genome | 30.8% | 58.1% | 51.1% | 41.7% |
| Reads mapped to exons | 25.6% | 46.8% | 43.2% | 33.7% |
| Reads mapped uniquely to genome | 29.8% | 52.6% | 49.8% | 38.5% |
| Estimated number of cells | 5,672 | 5,402 | 3,655 | 3,305 |
| Fraction of reads in cells | 94.40% | 94.50% | 93.2% | 93.3% |
| Median genes per cell | 2,239 | 3,360 | 3,199 | 3,011 |
| Total genes detected | 19,339 | 17,013 | 19,862 | 16,444 |

**Fig. 2.**
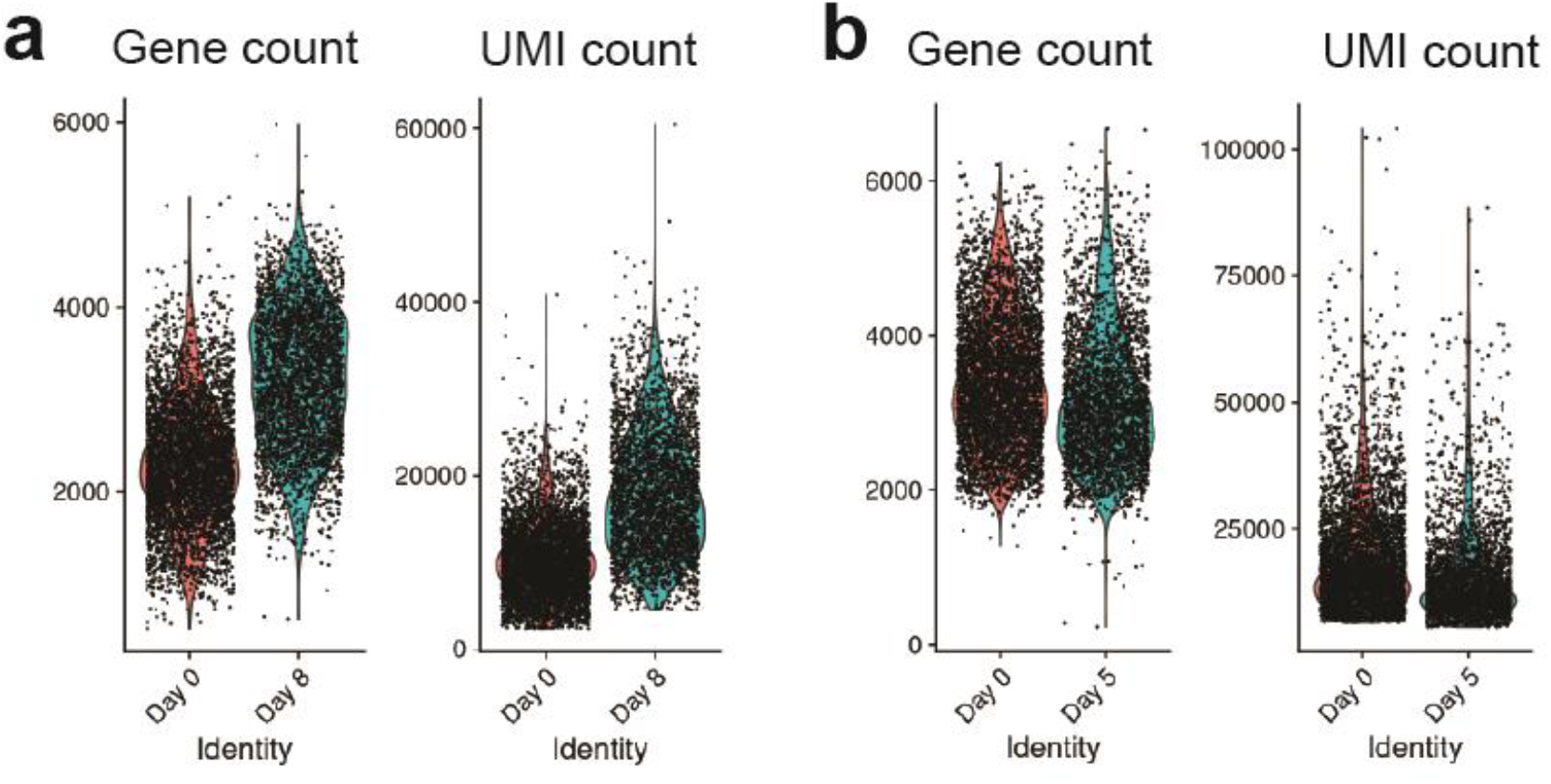
Single-cell RNA-Seq dataset quality assessment. **(a-b)** Violin plots of gene counts and UMI counts after quality control filtering in **(a)** SGBS cells and **(b)** 3T3-L1 cells, separated by the day of differentiation.

**Table 2.**
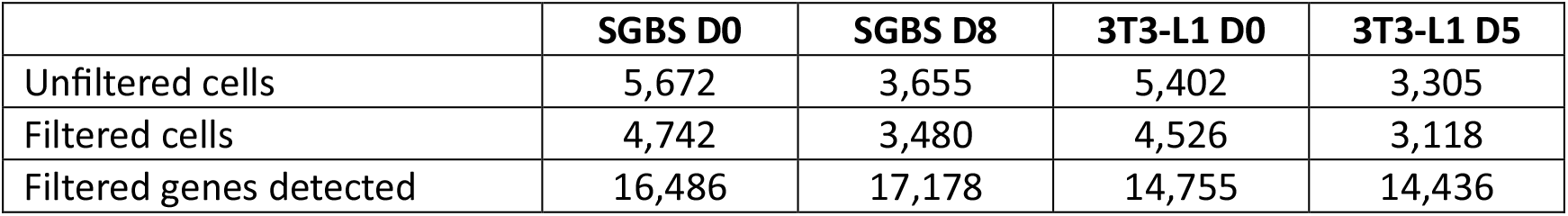
Final cell quantification statistics.

|  | SGBS D0 | SGBS D8 | 3T3-L1 D0 | 3T3-L1 D5 |
| --- | --- | --- | --- | --- |
| Unfiltered cells | 5,672 | 3,655 | 5,402 | 3,305 |
| Filtered cells | 4,742 | 3,480 | 4,526 | 3,118 |
| Filtered genes detected | 16,486 | 17,178 | 14,755 | 14,436 |

### Annotation of cell subpopulations

Adipogenesis is a highly heterogeneous process, and we expected the addition of differentiation stimuli to result in the appearance of additional cell states compared to D0 of differentiation, prior to the exposure to differentiation media. In fact, for both 3T3-L1 and SGBS cells we identified three cell clusters whose transcriptional profiles suggest they are preadipocytes, differentiating cells and adipocytes **(Fig. 3, Fig. 4)**. Furthermore, in both cell models there was a clear separation of cells isolated at D0, which corresponded to the preadipocyte clusters, and cells isolated after the induction of adipogenesis (D5 in 3T3-L1, D8 in SGBS), which corresponded to the other clusters **(Fig. 3, Fig. 4)**. Our scRNA-Seq dataset includes cells collected at two separate timepoints and processed independently, therefore we cannot rule out the presence of a batch effect contributing to the separation of D0 cells from later time points, which is a limitation of this study. However, analysis of the genes enriched in the identified cell clusters supports the view that the treatment with differentiation media affects the transcriptome, regardless of whether the cells fully differentiate, resulting in the differences between the clusters at D0 and D5/D8. In particular, adipogenesis is associated with major changes in the composition of the extracellular matrix (ECM) components. In line with previously published work, the preadipocyte cluster in SGBS cells showed enrichment in the expression of claudin 11 (*CLDN11*)*^16^*, and the clusters containing differentiating cells both in SGBS and 3T3-L1 models showed an enrichment of the expression of collagen type III alpha 1 chain (*COL3A1, Col3a1*) which is associated with adipogenic differentiation^17^. Furter, adipocyte markers fatty acid binding protein 4 (*FABP4*)*^18,19^*, adiponectin (*ADIPOQ*)*^20^*, and perilipin 4 (*PLIN4*)*^21^* were identified in the SGBS adipocyte cluster and *Fabp4^18,19^*, lipoprotein lipase (*Lpl*)*^22^*, and resistin (*Retn*)*^23^* were identified in the 3T3-L1 adipocyte cluster **(Table 3)**.

**Fig. 3.**
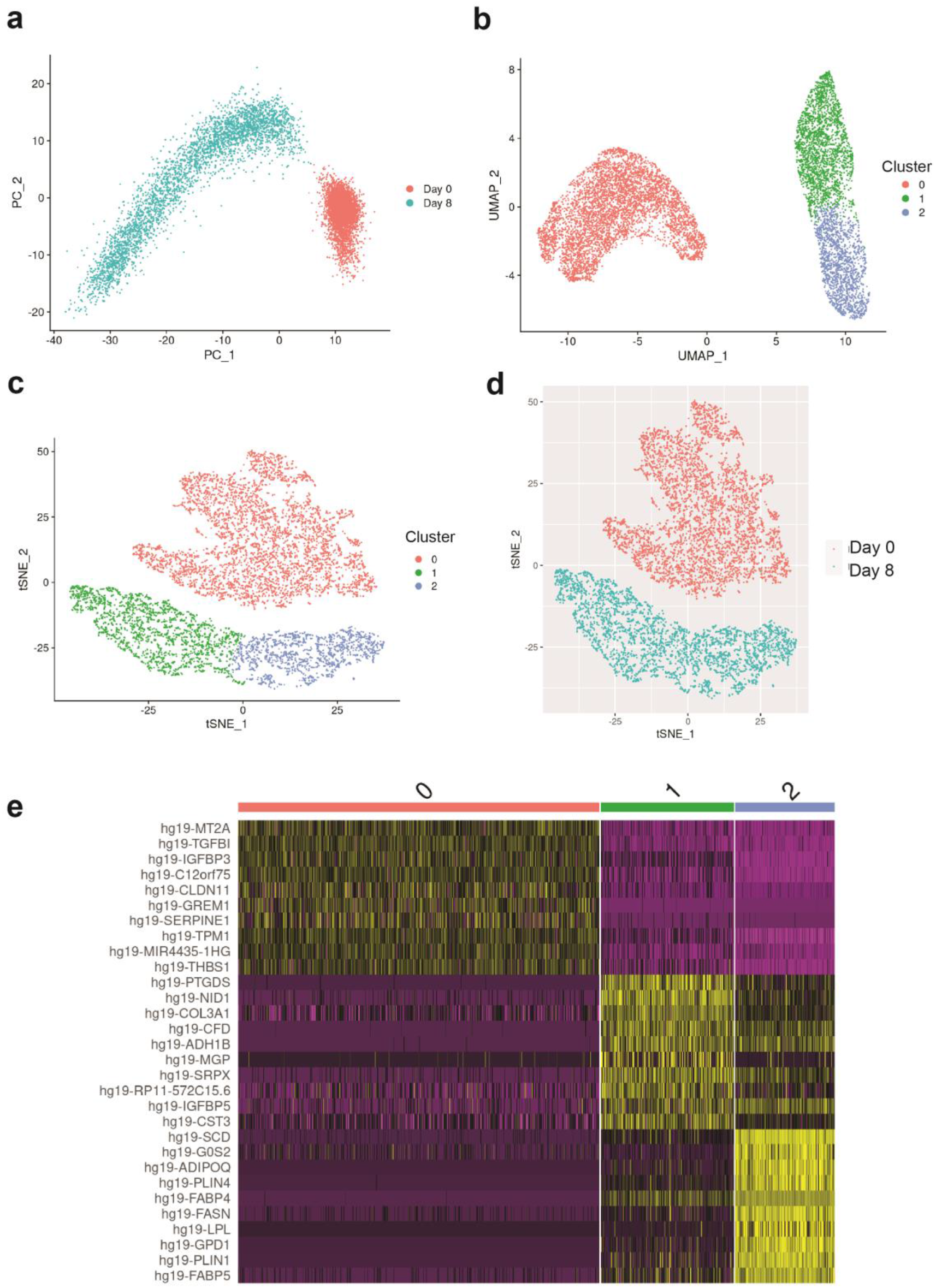
Clustering of scRNA-Seq data in human SGBS cells. **(a)** Primary component analysis (PCA) plot. **(b)** UMAP plot. **(c)** t-SNE plot. **(d)** Assignment of cells by differentiation day (D0 vs. D8), superimposed on the t-SNE plot. **(e)** Heatmap showing the expression of top 10 enriched genes per cell cluster.

**Fig. 4.**
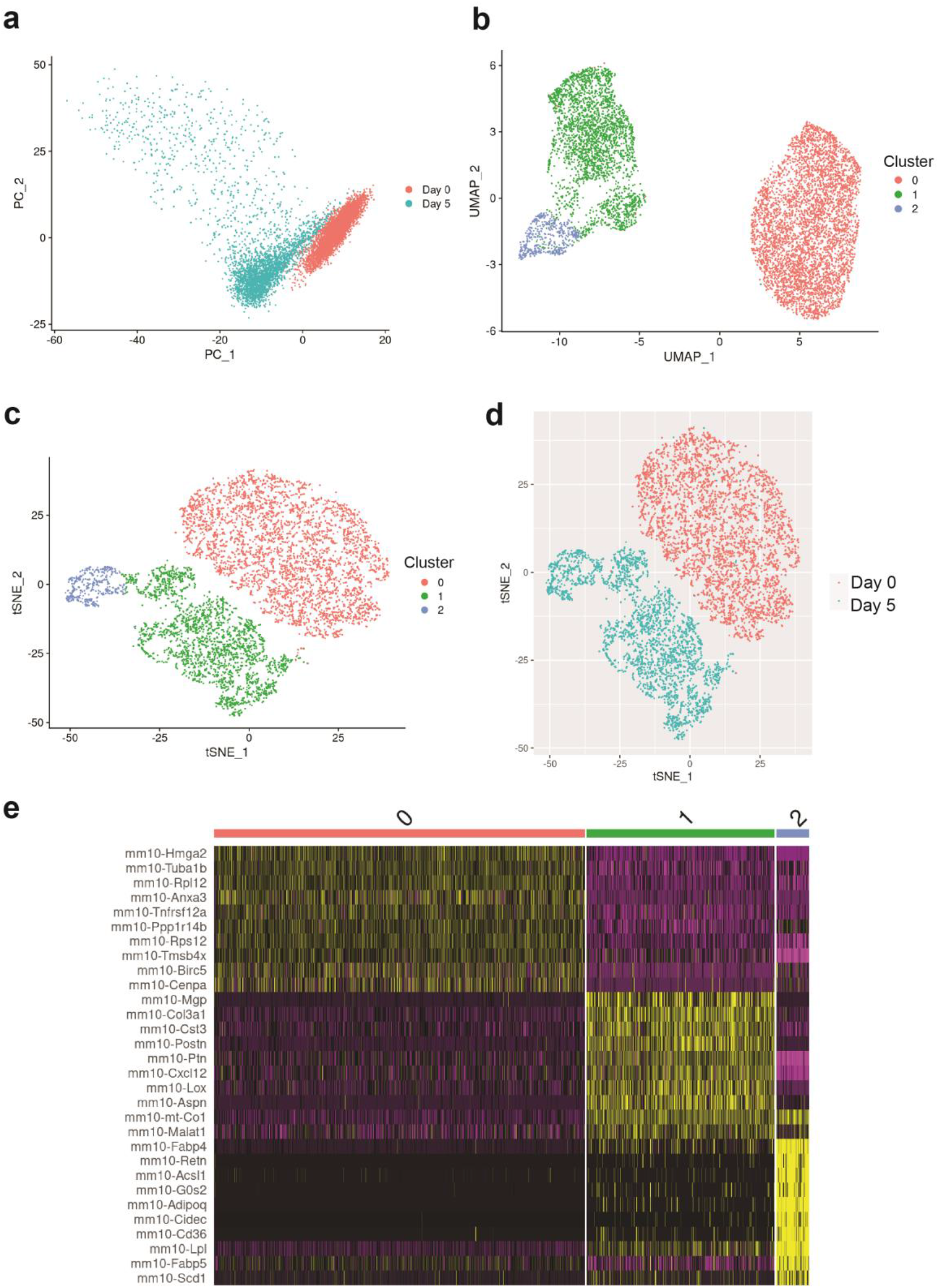
Clustering of scRNA-Seq data in murine 3T3-L1 cells. **(a)** Primary component analysis (PCA) plot. **(b)** UMAP plot. **(c)** t-SNE plot. **(d)** Assignment of cells by differentiation day (D0 vs. D5), superimposed on the t-SNE plot. **(e)** Heatmap showing the expression of top 10 enriched genes per cell cluster.

**Table 3.**
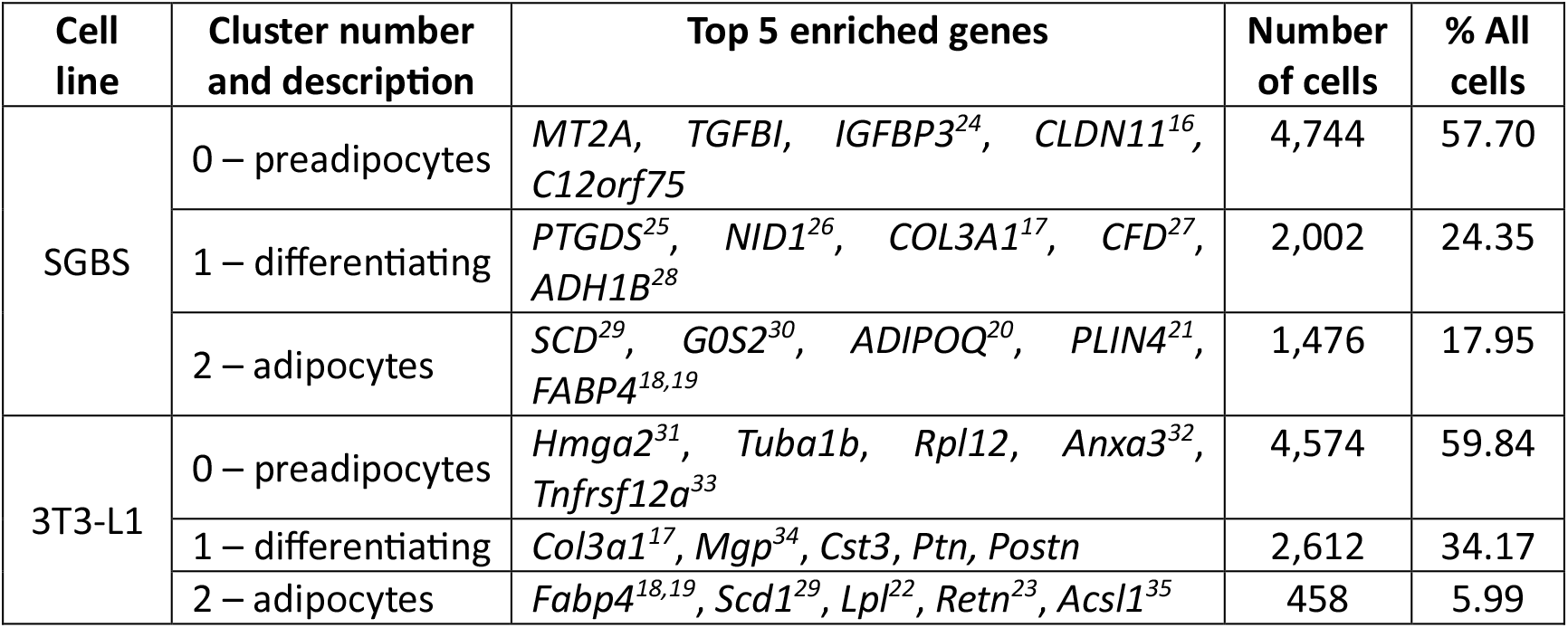
Description of cell clusters identified by unsupervised clustering.

## Code Availability

All analytical code used for processing and technical validation is available on the GitHub Repository (https://github.com/christopherjin/SGBS_3T3-L1_differentiation_scRNASeq). The provided R code was run and tested using R 4.2.2.

## Acknowledgements

The authors would like thank Dr. Erik Ingelsson for his support of this project. We acknowledge the technical assistance of the Stanford Genomics Service Center and the Stanford Shared FACS Facility.

E.B.M. was supported by the American Heart Association (AHA) postdoctoral fellowship (18POST34030448). T.Q. was supported by R01HL134817, R01HL139478, R01HL156846, R01HL151535, R01HL145708, UM1 HG011972 from the NIH, as well as by a Human Cell Atlas grant from the Chan Zuckerberg Foundation. J.W.K. was funded by NIH R01 DK116750, R01 DK120565, R01 DK106236, R01 DK107437, P30DK116074, and ADA 1-19-JDF-108.

## Author contributions

J.L. conceived the project, conducted experiments and analysed the data; C.J. conducted bioinformatic analyses, created figures and wrote the manuscript; S.G. assisted with bioinformatic analysis; A.R. assisted with bioinformatic analysis; M.W. provided critical resources for the project; C.Y.P. conceived the project and assisted with the experiments; T.Q. guided the bioinformatic analysis and critically reviewed the manuscript; E.B.M. analysed data, created figures and wrote the manuscript with the input from all the authors; J.W.K. oversaw the project and critically reviewed the manuscript.

